# Development of a Notch pathway assay and quantification of functional Notch pathway activity in T-cell acute lymphoblastic leukemia

**DOI:** 10.1101/2020.07.10.183731

**Authors:** Kirsten Canté-Barrett, Laurent Holtzer, Henk van Ooijen, Rico Hagelaar, Valentina Cordo, Wim Verhaegh, Anja van de Stolpe, Jules P.P. Meijerink

## Abstract

The Notch signal transduction pathway is pivotal for various physiological processes including immune responses, and has been implicated in the pathogenesis of many diseases including T-cell acute lymphoblastic leukemia. Various targeted drugs are available that inhibit Notch pathway signaling, but their effectiveness varies due to variable Notch pathway activity among individual patients. Quantitative measurement of Notch pathway activity is therefore essential to identify patients who could benefit from targeted treatment. We here describe a new assay that infers a quantitative Notch pathway activity score from mRNA levels of conserved direct NOTCH target genes. Following biological validation, we assessed Notch pathway activity in a cohort of TALL patient samples and related it to biological and clinical parameters including outcome. High Notch pathway activity was not limited to T-ALL samples harbouring strong *NOTCH1* mutations, including juxtamembrane domain mutations or hetero-dimerization combined with PEST-domain or *FBXW7* mutations, indicating that additional mechanisms may activate NOTCH signaling. The measured Notch pathway activity related to intracellular NOTCH levels, indicating that the pathway activity score more accurately reflects Notch pathway activity than predicted on the basis of *NOTCH1* mutations. Importantly, patients with low Notch pathway activity had a significantly shorter event-free survival compared to patients showing higher activity.

## Introduction

An increasing number of precision drugs is becoming available for clinical medicine, and many more are in development. These targeted drugs are intended for personalized medicine and aim at targeting pathophysiological defects underlying specific diseases in individual patients. For cancer, but also for many other diseases including auto-immune or immune-mediated diseases, patient samples may display a similar histopathology while significant pathophysiological variations can be found at the cellular level ^1, 2^. Such variations may be the reason that only part of all patients with a specific disease responds to a targeted drug. Matching the right drug to the right patient has therefore become an increasingly important issue. However, developing a diagnostic approach to reliably predict therapy responses has proven difficult. The prime example is oncology, where efforts in predicting patient responses to targeted drugs based on cancer genome mutations have generally been disappointing despite exceptions in select cases ^3–5 6, 7^. To improve clinical decision-making regarding targeted treatment and therefore to improve clinical outcome, assays are needed that accurately characterize and quantify the underlying pathophysiological processes in individual patient samples ^1, 8–17^. Cellular signal transduction pathways are evolutionary conserved and control fundamental cellular processes like cell division, differentiation, migration, and metabolism ^1, 18–20^. They include nuclear receptor pathways (*e.g*. androgen and estrogen receptor pathways), developmental pathways (Wnt, Hedgehog, TGFβ and Notch), the highly complex growth factor- and cytokine-regulated signaling pathway network including JAK-STAT, PI3K-AKT-mTOR and MAPK pathways, and the inflammatory NFκB pathway ^18, 21^. Measurement of the functional activity of these pathways in tumor biopsies from individual patients is expected to improve the prediction of therapy response. We have previously described a novel approach to quantitatively measure activity levels of individual signal transduction pathways in various cell and tissue types ^22–25^. In addition to development of assays to measure activity of the estrogen and androgen receptor pathways, the PI3K, JAK-STAT3, Wnt, Hedgehog, TGFβ, NFκB and JAK-STAT1/2 pathways, we now report the development and biological validation of a quantitative Notch pathway activity assay. The human Notch pathway is an evolutionary highly conserved developmental signaling pathway, activated by the interaction of one of four NOTCH transmembrane receptors with Jagged or Delta-like Canonical Notch ligands on neighboring cells ^16^. Upon ligand binding, the receptor is cleaved by two consecutive protease steps that include an ADAM (a disintegrin and metalloprotease domain containing) protease and the gamma-secretase complex. The resulting cleaved intracellular NOTCH (ICN) migrates to the nucleus where it forms a transcription factor complex with DNA binding factor RBPJ (recombination signal binding protein for immunoglobulin kappa J region) and coactivators of the MAML (Mastermind-like) family and activates transcription of its target genes. The Notch pathway plays a role in multiple diseases including T-cell acute lymphoblastic leukemia (T-ALL) ^16, 26^. Notch pathway inhibitors have been developed for multiple potential clinical applications, but their use has generally been associated with severe side effects ^27–32^. In addition, NOTCH inducers have been developed, *e.g*. for small cell lung cancer ^33^. A major clinical challenge is to minimize side effects and identify patients who benefit from Notch pathway modifying drugs.

To illustrate the potential utility of the Notch pathway assay for clinical decision-making, Notch pathway activity analysis was performed in a large cohort of diagnostic samples from pediatric T-ALL patients with a known genetic background and mutation status. Activating mutations in the NOTCH1 pathway including mutations in *NOTCH1* and/or *FBXW7—that* encodes for a ubiquitin ligase involved in the degradation of active intracellular NOTCH1 (ICN1)—are found in approximately 60% of T-ALL patients ^34, 35^. Publications on patient outcome in T-ALL report different prognostic significances for *NOTCH1*-activating mutations alone ^36^. We present evidence that patients with active Notch pathway signaling have a more favorable long-term outcome on high intensity combination treatment protocols ^37–39^.

## Methods

### Development of the Notch pathway assay

The mathematical approach to develop Bayesian network models for the measurement of signal transduction pathway activities based on mRNA expression analysis has been described in detail before ^24^. In brief, a causal computational network model for the Notch signal transduction pathway was generated that calculates the probability that NOTCH transcription factors are active based on the expression levels of direct target genes (**Fig. 1**). The Bayesian network describes the causal relation between up- or downregulation of NOTCH target genes and the presence of an active or inactive NOTCH transcription complex. Parameters that describe this relationship are based on literature evidence, and are calibrated on patient samples with known Notch pathway activity. Target genes for the Notch pathway assay were selected according to the same principles as described before, using available scientific literature ^22, 24^. The probesets of direct target genes from publicly available Affymetrix (Santa Clara, USA) HG-U133Plus2.0 microarray datasets were selected using the Bioconductor package hgu133plus2.db available in the statistical environment *R* and manually curated using GRCh38/hg38 available on the UCSC Genome Browser (www.genome.ucsc.edu, last access 2-17-2020) ^26, 28, 32, 40, 41^. Probesets representing intronic sequences, probesets on opposite strands or other chromosomal sequences than the respective target gene were excluded. Probesets that were missing in Bioconductor were added.

**Figure 1.**
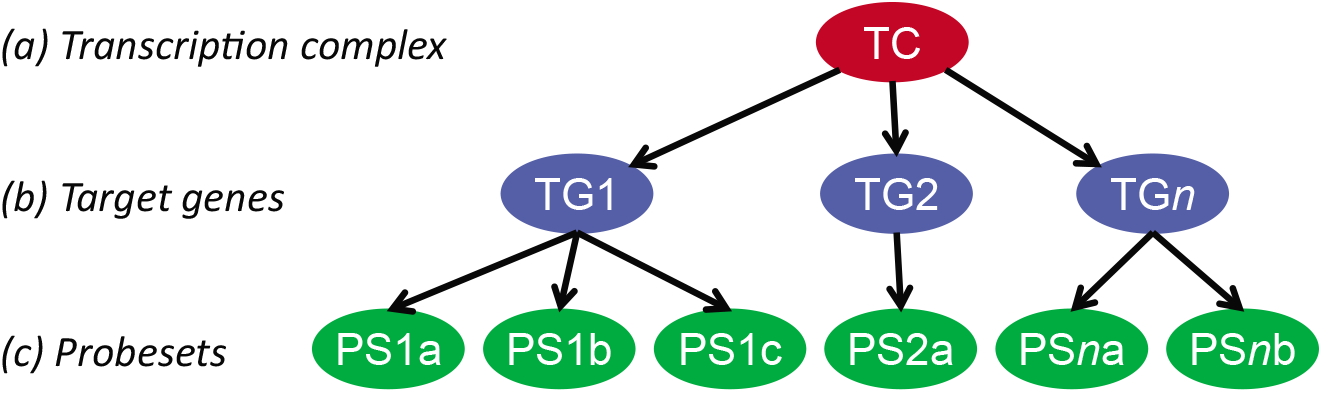
Bayesian model for the Notch signal transduction pathway. The structure of the Bayesian network used to model the transcriptional program of signaling pathways. The transcription complex refers to the transcription factor associated with a specific signal transduction pathway, which can be present in an inactive or in an active gene transcribing state; target genes refers to direct target genes of the transcription complex; probesets refer to probesets for the respective target gene present on Affymetrix HG-U133 Plus 2.0 microarray. With permission, ^24^.

### Calibration and validation of the Notch pathway activity model

The Notch pathway Bayesian model, intended for generic use across different cell and tissue types, was calibrated on a single public dataset containing data from normal (low Notch pathway activity) and high grade serous ovarian cancer (high Notch pathway activity) tissue samples ^42^. Following calibration, model parameters were frozen. Upon entering new mRNA probeset measurements into the model, Bayesian inference is used to calculate a linear scale log2 odds pathway activity score, as described ^22, 23^. The log2odds scale allows for high resolution detection of differences in signaling pathway activity. The model-based Notch pathway assay was validated using multiple independent Affymetrix datasets containing gene expression data from samples with known Notch pathway activity.

### Microarray data source and quality control

Affymetrix HG-U133Plus2.0 datasets used for Notch pathway model calibration, validation, and for Notch pathway analysis of T-ALL (GSE26713), are available at the GEO website (www.ncbi.nlm.nih.gov/geo, last access 2-17-2020). GEO datasets have been listed with associated publications in the figure legends. Before using the microarray data, extensive quality control was performed on Affymetrix data from each individual sample based on 12 different quality parameters according to Affymetrix’s recommendations and previously published literature ^22, 43, 44^ and then further preprocessed in the statistical environment *R* using frozen RMA ^45^ with ‘robust weighted average’ summarization.

### Description of the T-ALL pediatric patient cohort

Affymetrix HG-U133Plus2.0 gene expression profiles (GSE26713) from diagnostic biopsies of 117 T-ALL patients who were treated according to the German co-operative study group for childhood ALL-97 protocol (COALL-97) or the Dutch Childhood Oncology Group (DCOG) protocols ALL-7,-8 or −9 were used in this study ^46^. The patient data used in this study was obtained with informed consent from the subjects’ guardians and in accordance with the Declaration of Helsinki.

### Statistics

For the validations of the Notch pathway model, two-sided Wilcoxon signed-rank statistical tests were performed. Other used statistical methods that are more appropriate due to the content of a specific dataset are indicated in figure legends. For pathway correlation statistics, both Pearson correlation and Spearman rank correlation tests were performed; since the results were similar, only the Pearson correlation coefficient and associated p-value are reported. For outcome analysis, Kaplan-Meier survival curves were calculated together with associated p-value using the log-rank test.

## Results

### Development of the Notch pathway assay and selection of NOTCH target genes

For the development of the Notch pathway assay, we selected high evidence direct target genes of NOTCH. This selection is based on (i) the presence of minimally one binding element in the promoter region, (ii) functionality of these binding elements that have been assessed for instance by gene promoter-reporter studies, (iii) binding of ICN to the respective response/enhancer element using ChIP and/or Electrophoretic Mobility Shift Assay, (iv) their differential expression upon pathway activation and/or inhibition, and (v) consistency of evidence as reported by multiple research groups for multiple cell/tissue types. Based on such accumulated experimental evidence as described before ^22–24^, we selected 18 direct target genes *CD44, DTX1, EPHB3, HES1, HES4, HES5, HES7, HEY1, HEY2, HEYL, MYC, NFKB2, NOX1, NRARP, PBX1, PIN1, PLXND1, SOX9* (**Supplemental Table S1**)^47–89^. This number is sufficient for robust and sensitive prediction of the pathway activity while comprising only high evidence target genes that enable maximal specificity over multiple cell types.

### Calibration and validation of the Notch pathway activity assay

We calibrated the Notch pathway assay using data from high-grade serous (HGS) ovarian cancer samples with high Notch pathway activity and normal ovarian tissue samples with low Notch pathway activity (**Fig. 2A**). While in healthy ovarian tissue samples the Notch pathway is inactive, HGS ovarian cancer is associated with an active Notch pathway and activating *NOTCH3* gene mutations or amplifications in about two third of the patients ^42, 90–92^. Following freezing of the Notch pathway model, it was validated on various independent datasets from cells of different tissue origins with Notch pathway-activated or gamma-secretase inhibited conditions, including cell types from ectodermal (neuroblastoma) and endodermal (lung cancer cells) origin in contrast to the mesodermal origin of the ovarian cancer samples the model had been calibrated on (**Figs. 2B-I**). Two independent clones of a neuroblastoma cell line transfected with ICN3 show a rapid and persistent quantitative increase in Notch pathway activity score starting within 4 hrs and reaching a plateau activity at 12 hours after transfection (**Fig. 2B**). In leukemia, the AF1Q-MLLT11 fusion protein confers sensitivity to ligand-induced Notch pathway signaling ^93, 94^. Hematopoietic progenitor cells (CD34^+^CD45RA^-^Lin^-^) from umbilical cord blood were transduced with the A2M mutant version of this fusion product that sequesters it in the nucleus. Following a three-day exposure to immobilized NOTCH ligand Delta1ext-IgG at two different dose levels, high Notch pathway activity scores were measured for both mock and A2M transduced cells (**Fig. 2C**). As expected, Notch pathway activity scores are higher for A2M-transduced cells than control cells.

**Figure 2.**
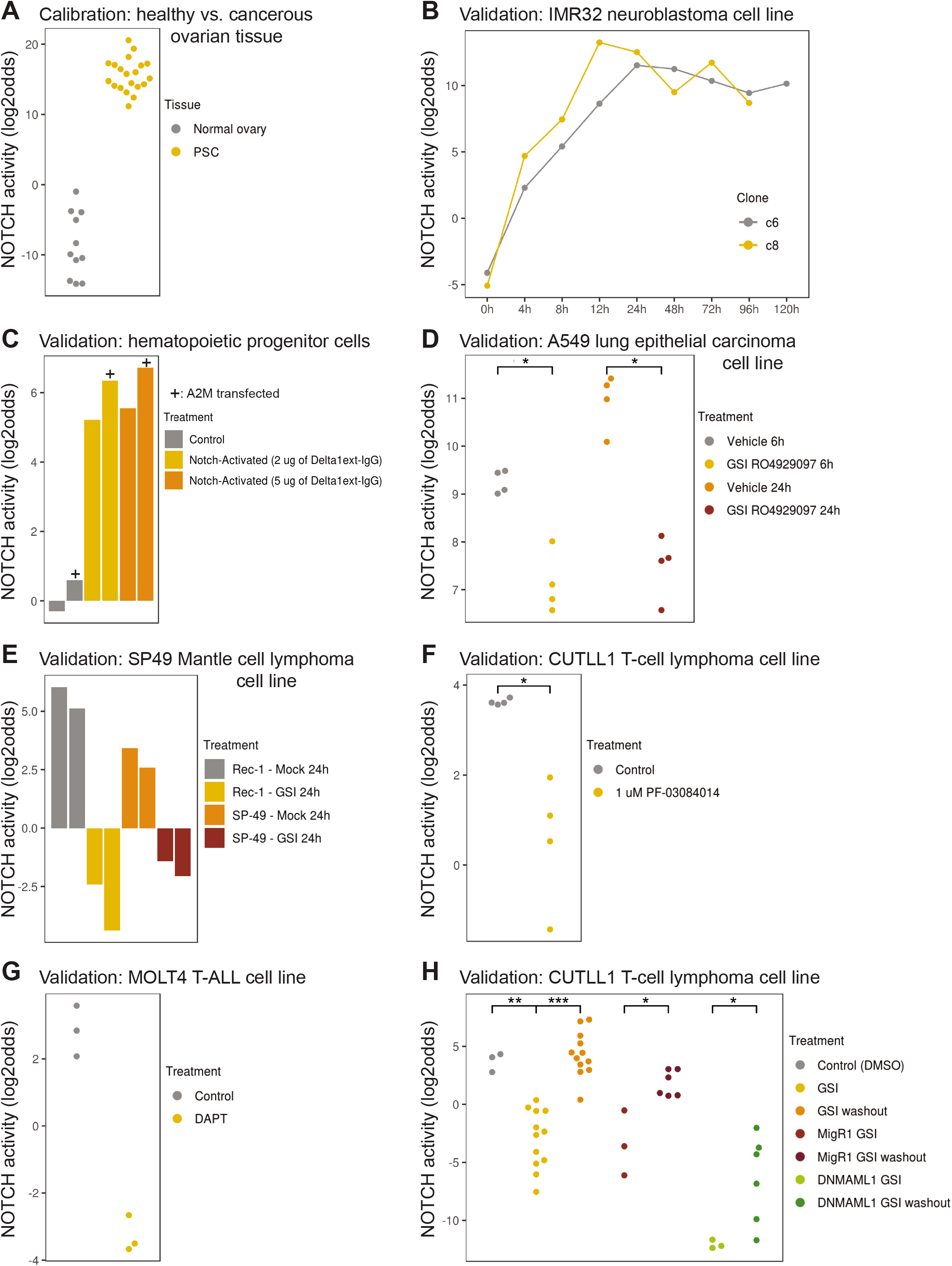
Calibration and biological validation of the Notch pathway model. (**A**) Calibration of the Notch pathway model. GSE7307, GSE18520 ^105^, GSE29450 ^106^, GSE36668 ^107^, normal ovarian tissue samples (inactive pathway); GSE2109, GSE9891 ^108^, high grade serous ovarian cancer samples (active pathway). (**B-G**) Validation of the model on independent GEO datasets from different cell lines. *p<0.05, **p<0.01, ***p<0.001. (**B**) GSE16477.^109^ Two clones (c6 and c8) of the IMR32 neuroblastoma cell line at different times (0-120 hours) after induction of active intracellular NOTCH3. (**C**) GSE29524. A2M (+ symbol) or control vector transfected CD34^+^CD45RA^-^Lin^-^ hematopoietic progenitor cells from umbilical cord blood were cultured for 72hrs on a surface with 0, 2 or 5μg plastic-immobilized NOTCH ligand Delta1ext-IgG. A2M is a nuclear-trapped mutant of AF1q/MLLT11. (**D**) GSE36176.^110^ A549 lung cancer cell line subjected to vehicle control or gamma secretase inhibitor (GSI) RO4929097 for 6 or 24 hours. (**E**) GSE34602.^95^ Rec-1 (containing an activating *NOTCH1* mutation) and SP49 Mantle cell lymphoma cell lines subjected to vehicle control or GSI compound E for 24hrs. SP49 cells harbor an activating NOTCH4 rearrangement. (**F**) GSE33562.^111^ Duplicate samples of the CUTLL1 T-cell lymphoma cell line were subjected to vehicle control or the GSI PF-03084014 (1μM) for 48hrs. (**G**) GSE6495.^112^ MOLT4 T-cell acute lymphoblastic leukemia cell line before and 48 hours after addition of the GSI DAPT (5μM); three independent experiments. (**H**) GSE29544.^113^ CUTLL1 T-cell lymphoblastic lymphoma cells subjected to GSI compound E (1μM) for 3 days. From left to right: DMSO control; 3 grouped conditions: GSI without or with 2 or 4 hours mock washout; 4 grouped conditions: GSI followed by 2 or 4 hours GSI washout in the presence or absence of cycloheximide; GSI in presence of a control viral transcript MigR1; 2 grouped conditions: GSI in the presence of a control viral transcript MigR1 with 2 or 4 hours washout; GSI in the presence of viral transcript DNMAML1; 2 grouped conditions: GSI in the presence of viral transcript DNMAML1 with 2 or 4 hours washout. The activity score is calculated as log2odds. Two-sided Wilcoxon signed-rank statistical tests were performed, p-values are indicated in de figures. In case fewer than 3 samples were needed for presentation, bar plots are used instead of dot plots.

This result provides additional evidence for the ability of the Notch pathway assay to quantify small differences in Notch pathway activity. In other experimental designs in which Notch pathway activity was inhibited by exposure to gamma-secretase inhibitors (GSIs), the robustness of the assay in various additional cell types was validated. A549 lung cancer cells exposed to the GSI RO4929097 for 6 or 24 hours scored a significantly lower Notch pathway activity than control A549 cells (**Fig. 2D**). Similar findings were found for the GSI-exposed Mantle B-cell lymphoma cell line SP-49 and the NOTCH mutant Rec-1 line ^95^ (**Fig. 2E**), and for the T-cell lymphoma and leukemia cell lines CUTLL1 and MOLT4 (**Figs. 2F,G**). In CUTLL1 cells, wash-out of the GSI resulted in reactivation the Notch pathway, which was accurately quantified (**Fig. 2H**). Furthermore, a dominant-negative form of the NOTCH cofactor MAML1 (DNMAML1) synergized with GSI and resulted in the lowest Notch pathway activity score. Interestingly, in this study the removal of GSI was performed both in the absence or presence of the protein translation inhibitor cycloheximide to exclude any feedback or secondary effects from NOTCH induced gene products. The measured Notch pathway activity scores were independent of protein translation, confirming that all genes that are part of the computational Notch pathway model are indeed direct target genes (**Supplemental Fig. S1**). In summary, these results demonstrate that the ovarian cancer-calibrated Notch pathway assay can be used to measure Notch pathway activity levels in T-cells, while the limited results available on other cell types suggest that the assay may also be usable in cell types of endodermal and ectodermal origin.

### Notch pathway activities in pediatric T-ALL patient samples

Following biological validation of the Notch pathway assay, we measured Notch pathway activity scores in diagnostic samples from 117 pediatric T-ALL patients. This dataset has been previously used to distinguish four main T-ALL subgroups (ETP-ALL/immature, TLX, Proliferative and TALLMO) based on their differential gene expression profiles that strongly correlate with unique oncogenic rearrangements ^46^. Notch pathway activity scores ranged from −8.59 to 7.45 on the linear log2 odds scale. To investigate these scores in relation to the presence of specific types of Notch pathway-activating mutations, we categorized NOTCH1 mutations into weak or strong activating mutations as done before ^35, 96, 97^: weak NOTCH1 activating mutations are considered mutations in the NOTCH1 heterodimerization domain (HD), PEST domain, or inactivating mutations in *FBXW7*. Strong NOTCH1-activating mutations are mutations in the juxtamembrane domain or HD-mutations combined with PEST domain or *FBXW7* mutations. Based on this division, the median Notch pathway activity score was lowest for the patient samples without NOTCH-activating mutations and highest for the samples with the strong NOTCH1-activating mutations (p<0.001; **Fig. 3A**). Still, there is considerable overlap in activity scores among these groups. To investigate a potential effect of differences in genetic backgrounds among patients, we compared Notch pathway activity levels between the four T-ALL subtypes. The TLX subtype had the highest Notch pathway activity scores compared to the other subtypes and included 10 out of 23 patient samples with strong NOTCH mutations (**Fig. 3B**). Various TLX samples without or with only weak NOTCH-activating mutations also had high Notch pathway activity scores, further supporting the previous observation that alternative Notch pathway-activating mechanisms may exist. We then related activity scores to intracellular NOTCH1 (ICN1) levels as measured using reverse-phase protein array for 62 patient samples ^35^. We observed a significant relationship between ICN1 levels and the absence or presence of NOTCH1-activating mutations (**Fig. 3C**) and between ICN1 levels and the Notch pathway activity scores (**Fig. 3D**). The significance of the correlation between ICN1 levels and Notch pathway activity was mainly attributed to the strong NOTCH1-activating mutations, as the significance was lost for patient samples without or with only weak NOTCH1-activating mutations (**Supplemental Fig. S2**). This raised the question whether those samples could harbor other Notch pathway-activating mechanisms. For this, we assessed NOTCH3 protein levels as an alternative Notch pathway-activating mechanism for various *NOTCH1/FBXW7* non-mutated T-ALL patient samples with low ICN1 levels but high Notch pathway activity scores. We did not find expression of NOTCH3 protein in these and other T-ALL samples tested (not shown). We then excluded an influence of bone marrow or peripheral blood origin of the T-ALL samples on Notch pathway activity scores (not shown). Therefore, the incidental discrepancy between ICN and Notch pathway activity scores remains unclear. In conclusion, the results show that the Notch pathway assay quantitatively measures Notch pathway activity not only in cell line systems, but also in a cohort of primary T-ALL patient samples.

**Figure 3.**
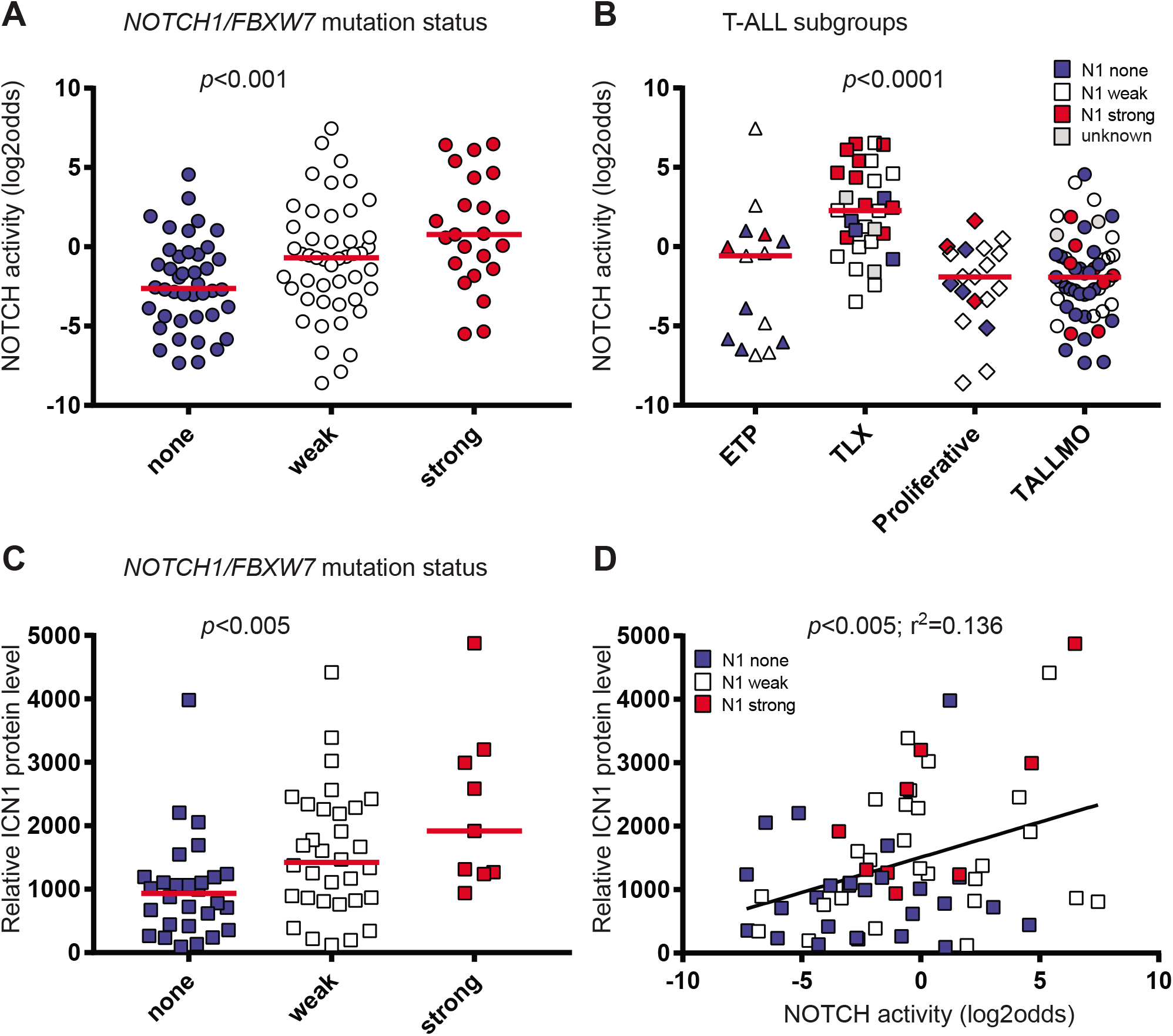
Notch pathway activity in T-ALL. (**A-D**) GSE26713.^46^ No NOTCH activating mutations (blue symbols), weak NOTCH1 activating mutations (NOTCH1 heterodimerization domain, PEST domain or in FBXW7) (white symbols) and strong NOTCH1 activating mutations (juxtamembrane domain or more than one NOTCH1 activating mutation) (red symbols) are indicated. P-values are indicated. (**A-C**) Kruskal-Wallis statistical test. Medians are indicated by the red lines. (**D**) Linear regression test. (**A**) Notch pathway activity of T-ALL samples (n=112) per *NOTCH1/FBXW7* mutation status group. (**B**) Notch pathway activity per T-ALL subgroup (n=117). Five samples have an unknown *NOTCH1/FBXW7* mutation status (grey symbols). (**C**) Active intracellular NOTCH1 (ICN1) protein level measured in relative intensity units using reverse-phase protein array (RPPA), indicated per *NOTCH1/FBXW7* mutation status group (n=69). (**D**) Correlation of active intracellular NOTCH1 (ICN1) protein level and Notch pathway activity (n=62).

### Notch pathway activity and T-ALL patient survival

The prognostic significance of NOTCH-activating mutations is not consistent in various patient studies ^36^. Part of this may be due to mechanisms, other than activating mutations in hotspots of *NOTCH1* or *FBXW7*, that activate NOTCH signaling in T-ALL patients, and may explain the large overlap in Notch pathway activity levels for T-ALL patients with and without *NOTCH/FBXW7* mutations. In order to investigate outcome in relation to Notch pathway activity, we divided the T-ALL patients into three groups based on their NOTCH activity scores: a group with the highest NOTCH activity scores (>75^th^ percentile), a group with the lowest activity scores (<25^th^ percentile) and a group with intermediate activity scores (between the 25^th^-75^th^ percentiles of activity scores). When assessing the event-free and relapse-free survival curves, we observed that the patients with lowest activity scores had the shortest event-free survival compared to both other groups (p<0.05), while relapse-free survival showed the same trend (**Figs. 4A,B**).

**Figure 4.**
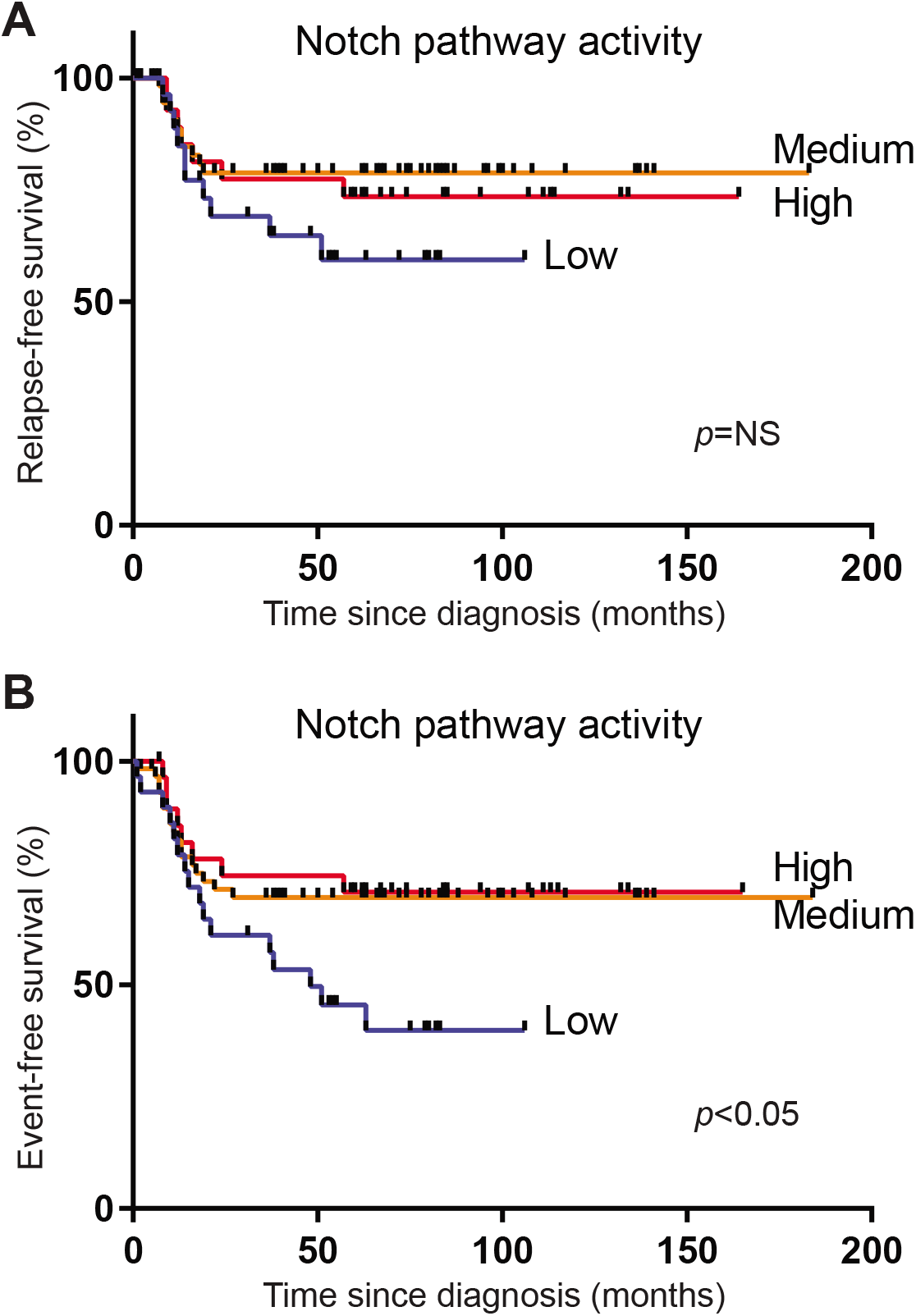
Relapse-free and event-free survival of T-ALL patients in three different Notch pathway activity groups. Three different Notch pathway activity groups were separated based on the lowest 25% Notch pathway activity (blue line), the highest 25% Notch pathway activity (red line), and the remaining 50% termed ‘middle’ Notch pathway activity (orange line). Relapse-free (**A**) and event-free (**B**) survival is plotted for T-ALL pediatric patients treated on the DCOG ALL-7, −8, and −9 and COALL-97 protocols. p=NS (not significant) (**A**) and p<0.05 (**B**) (log-rank test). Events include relapse, non-responsiveness to induction or maintenance therapy, change of treatment or death due to infection, toxicity or other causes.

### Relation between Notch pathway activity and PTEN loss

The group with the lowest NOTCH activity scores contained patients that lacked either PTEN protein and/or had inactivating mutations or deletions in *PTEN*. We found an increased percentage of patients (11 out of 29, 38%) with functional PTEN loss in the group with the lowest Notch pathway activity, whereas only 12 out of 84 patients (14%) with intermediate and high Notch pathway activity scores had functional PTEN loss (p=0.006, Pearson Chi-Square, 2-sided).

## Discussion

We have developed an assay to measure Notch pathway activity, consisting of a Bayesian network computational model which calculates a pathway activity score based on target gene expression levels. The set of NOTCH target genes was selected based on experimental evidence, irrespective of cell type or gene function ^22–24^. The computational model was successfully validated on a variety of samples from different cell types with known Notch pathway activity, *i.e*. brain, lung, hematopoietic stem cells, and T-ALL cell lines. This suggests that the assay can be used on multiple different cell types without model recalibration, even across cell types originating from different embryonic germ layers. This is to a large extent enabled by the selection of high evidence *direct* transcriptional target genes of the NOTCH transcription factor family (*e.g*. NOTCH1, NOTCH2, NOTCH3), eliminating cell type-specific influences on target gene expression as much as possible. In addition, the Bayesian network model is well suited to handle variations in input data, which presents a crucial advantage when analyzing patient samples that are intrinsically highly variable in gene expression regulation ^22^. Other RNA-based pathway analysis tools are available, mainly for biomarker discovery applications, and differences have been discussed before ^22, 24, 98–100^. In short, we use a knowledge-based Bayesian modelling approach as opposed to a more generally used data-driven approach, thus avoiding common problems with data-overfitting. This approach improves specificity in measuring signaling pathway activity, and enables development as a diagnostic assay across multiple disease types.

To explore clinical utility of the biologically validated Notch pathway model, we have analyzed diagnostic samples of 117 pediatric T-ALL patients. We found that the Notch pathway activity score was related to the presence of NOTCH1-activating mutations and the type of mutations, and correlated to the levels of ICN protein in these samples. Correspondingly, we found the highest Notch pathway activity scores in the TLX subgroup, a group that we described before to have the highest incidence of NOTCH1-activating mutations ^35^. Most T-ALL patients in this T-ALL subgroup (21 out of 30 patients) bear *TLX3-BCL11B* rearrangements ^46^. Moreover, the TLX subgroup is related to gamma-delta T-cell lineage development ^101^. Interestingly, human gammadelta T-cell lineage development especially depends on high Notch pathway activity levels, in contrast to alpha-beta T-lineage development ^84^. The proliferative and TALLMO subgroups that are associated with early and late cortical stages of the alpha-beta T-lineage, respectively, indeed have lower Notch pathway activity scores. Therefore, the NOTCH dependency in normal development mirrors that of the respective T-ALL subgroups. Remarkably, about half of the ETP-ALL patients seem to have an activated Notch signaling pathway based on measured activity scores, despite their overall lower incidence of NOTCH-activating mutations ^102^. We observed that various samples without or with weak NOTCH-activating mutations still have high Notch pathway activity scores ^35^. This is especially evident for patients from the TLX subgroup and points to other, yet unidentified, mutations outside the present hotspot regions or other mechanisms that may activate the Notch pathway in T-ALL.

Patients with a Notch pathway activity score in the lowest 25^th^ percentile had the worst event-free and relapse-free survival. Interestingly, *NOTCH* mutations in this cohort were not associated with beneficial outcome as reported before ^35^ while other studies identified activating *NOTCH* mutations as a favorable prognostic factor ^37–39^. This result suggests that scoring the Notch pathway activity might be a more reliable method to determine prognosis than identifying NOTCH-activating mutations. In addition, the Notch pathway test has the potential to improve stratification of patients to novel therapies targeting the Notch pathway.

Patients with the lowest Notch pathway activity scores were more likely to have functional PTEN loss, indicating that Notch pathway activation and PTEN inactivation reflect two distinct T-ALL entities as we and others reported before ^97, 103^. PTEN aberrancies are often found in the TALLMO T-ALL subgroup in which they occur mutually exclusive with strong NOTCH1 mutations ^97^.

Moreover, patients with PTEN aberrancies have been shown to have an inferior survival ^97, 103^. The finding that PTEN aberrancies occurred more often in the patients with the lowest Notch pathway activity helps explain the inferior event-free/relapse-free survival of this group.

Overall, our results indicate that Notch pathway activity cannot be deduced from the presence of activating mutations only, which may provide an explanation for the differences in the prognostic significance of NOTCH-activating mutations in various pediatric and adult patient cohorts ^36^. While the here described Notch pathway assay is expected to be of value for a broad range of diseases as well as for preclinical research and drug development, the first envisioned clinical application is therapy response prediction, *e.g*. to NOTCH inhibitors, for T-ALL, small cell lung cancer, and other malignancies. To enable use of the Notch pathway activity assay on formalin fixed paraffin embedded tissue samples that are the standard in pathology diagnostics, the here described Affymetrix-based Notch pathway activity test has been converted to an RT-qPCR based test; to enable determination of Notch pathway activity on RNA sequencing data, the assay has been converted to an RNAseq-based assay (www.philips.com/oncosignal). The conversion procedure has been described before, and does not involve addition of new target genes ^104^.These assays will be used in future clinical validation studies.

## Author contributions

KCB performed T-ALL-related experiments, analyzed patient data and wrote manuscript, LH developed and validated the Notch pathway assay, wrote manuscript, HvO developed the Notch pathway assay, RH performed statistical analyses using patient data, VC performed experiments, WV developed the Notch pathway assay, AvdS developed and validated the Notch pathway assay, wrote manuscript, JPPM explored the clinical utility of the Notch pathway assay using T-ALL patient samples, analyzed patient data, wrote manuscript.

## Acknowledgements

KCB and RH are funded by the Dutch ‘Kinderen Kanker Vrij’ foundation grants KiKa-295 and KiKa-219, respectively. VC is funded by the Dutch Cancer Society grant KWF-10355. We would like to thank Sieglinde Neerken for detailed reading of the manuscript and providing valuable comments.

