## Supplement for "Development of a Notch pathway assay and quantification of functional Notch pathway activity in T-cell acute lymphoblastic leukemia"

### Supplemental Figure legends:

**Supplemental Figure S1. Extended validation of the Notch pathway model in CUTLL1 cells.**
Related to Fig. 2H., dataset GSE29544. CUTLL1 cells subjected to GSI compound E (1µM) for 3 days.
(**A**) From left to right: DMSO control; GSI; GSI followed by 2 hours washout of the GSI; GSI followed by 2 hours washout of the GSI in the presence of cycloheximide (CHX); GSI followed by 4 hours washout of the GSI; GSI followed by 4 hours washout of the GSI in the presence of CHX; GSI with 2 hours mock washout; GSI with 4 hours mock washout.
(**B**) From left to right: DMSO control; GSI in presence of a control viral transcript (MigR1); GSI in the presence of MigR1 with 2 hours washout; GSI in the presence of MigR1 with 4 hours washout; GSI in the presence of a dominant negative viral transcript DNMAML1; GSI in the presence of DNMAML1 with 2 hours washout; GSI in the presence of DNMAML1 with 4 hours washout.

**Supplemental Figure S2. Correlations of ICN1 level and Notch pathway activity, divided per NOTCH1-activating mutation status.**
Related to Fig. 3D. No NOTCH activating mutations (blue symbols), weak NOTCH1 activating mutations (NOTCH1 heterodimerization domain, PEST domain or in FBXW7) (white symbols) and strong NOTCH1 activating mutations (juxtamembrane domain or more than one NOTCH1 activating mutation) (red symbols) are indicated. P-values are indicated. NS: not significant. Correlation of ICN1 protein level and NOTCH pathway activity for samples without and with weak NOTCH1-activating mutations (**A**), with strong NOTCH1-activating mutations (**B**), without NOTCH1-activating mutations (**C**), and with weak NOTCH1-activating mutations (**D**).

**Supplemental Table S1. Scoring of NOTCH target genes with respect to evidence as to direct gene regulation by Notch transcription factors.** Evidence score calculation: Response element (1); ChIP (1); Differential expression (down), *e.g.* by GSI treatment (1); Differential expression (up), *e.g.* by ICN1 transfection (1); Luciferase assay (1); Cycloheximide addition (1); EMSA (1); Evidence obtained in mouse (0.5 instead of 1).

| # | Gene | Evidence + references | Evidence score |
|---|---|---|---|
| 1 | *CD44* | Response element [78], ChIP [78], GSI treatment [61, 78] | 3 |
| 2 | *DTX1* | GSI treatment [73, 83, 86], Notch1 knockdown [73], Notch1 induction (mouse) [50] | 1.5 |
| 3 | *EPHB3* | Response element [55, 78], ChIP [55, 78], GSI treatment [55, 78], Luciferase (Depending on cell line) [55], dnMAML induction [55] | 4 |
| 4 | *HES1* | Response element [57, 78], ChIP [55, 78], GSI treatment [72, 78, 79, 82, 86], dnMAML induction [55], Luciferase [80], Luciferase (mouse) [56], Delta1+CHX (mouse) [62] | 4.5 |
| 5 | *HES4* | Response element [85], ChIP [82], GSI treatment [72, 82] | 3 |
| 6 | *HES5* | Response element [57], Notch mutant (mouse) [49], GSI treatment [64, 82], NICD transfection (mouse) [53] | 2.5 |
| 7 | *HES7* | GSI treatment [51], Response element (mouse) [47], Luciferase assay (mouse) [47] | 2 |
| 8 | *HEY1* | Response element [52, 57], Luciferase (mouse) [68], Cycloheximide (mouse) [54], ChIP [52], GSI treatment [46, 82, 86] | 4 |
| 9 | *HEY2* | Response element [57], Luciferase (mouse) [53, 68], Promoter deletions (mouse) [68], GSI treatment [82], N4ICD overexpression [76], CSL mutation [76], immobilized Dll1 [58] N1ICD transfection (mouse) [53, 65], ChIP (mouse) [65], ChIP (human) [60], EMSA [60] | 5.5 |
| 10 | *HEYL* | Response element [57, 68], GSI treatment [9], Notch1 knockout (mice) [63], Notch activitation(mouse) [68], Promoter deletion (mouse) [68] | 2.5 |
| 11 | *MYC* | Response element [86], GSI treatment [72, 73, 82, 86], GSI treatment (mouse) [81], Cycloheximide treatment [86], ChIP [66, 73, 86], ChIP (mouse) [81], EMSA [86], Notch1 knockdown [73] | 5 |
| 12 | *NFKB2* | Response element [71], EMSA [71], luciferase [71], ChIP [84] | 4 |
| 13 | *NOX1* | Response element [78], ChIP [78] , GSI treatment [78] | 3 |
| 14 | *NRARP* | Response element [75], GSI treatment [82, 83], Luciferase (mouse) [75], EMSA [75], NICD Mutation (mouse) [59], N3ICD transfection [87] | 4.5 |
| 15 | *PBX1* | Response element [74], Cycloheximide treatment [74], GSI treatment [74], N3ICD inhibition [67, 74] | 4 |
| 16 | *PIN1* | Response element [79], Luciferase assay [79], ChIP [48, 79], GSI treatment [79], N1ICD overexpression [79] | 5 |
| 17 | *PLXND1* | Response element [77], Luciferase assay [77], dnRBPj [77], N1ICD overexpression [77] | 4 |
| 18 | *SOX9* | Response element [78, 88], ChIP [78, 88], GSI treatment [78], Notch1 signaling induction [69, 70], Cycloheximide treatment [69, 70] | 5 |

**Supplemental Figure S1. Extended validation of the Notch pathway model in CUTLL1 cells.**

A

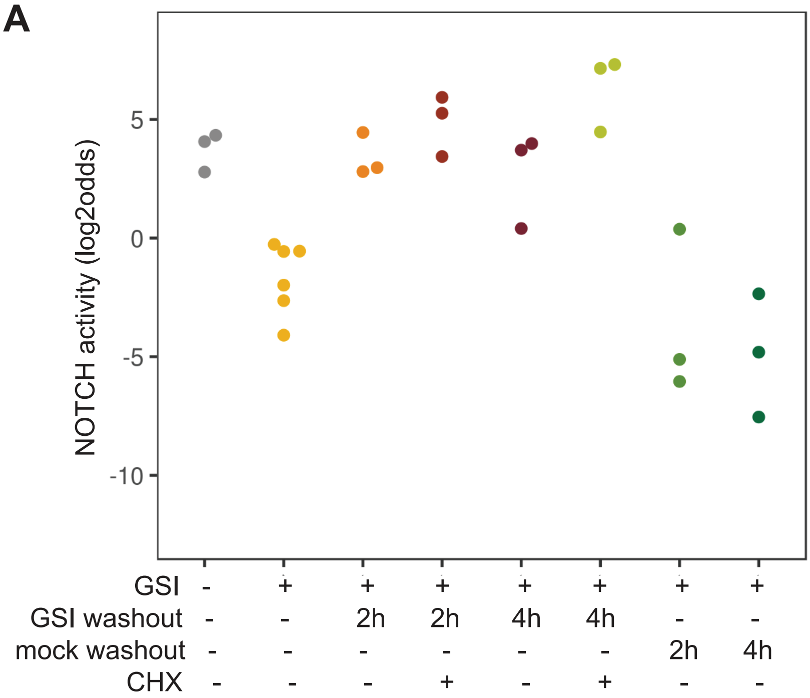


B

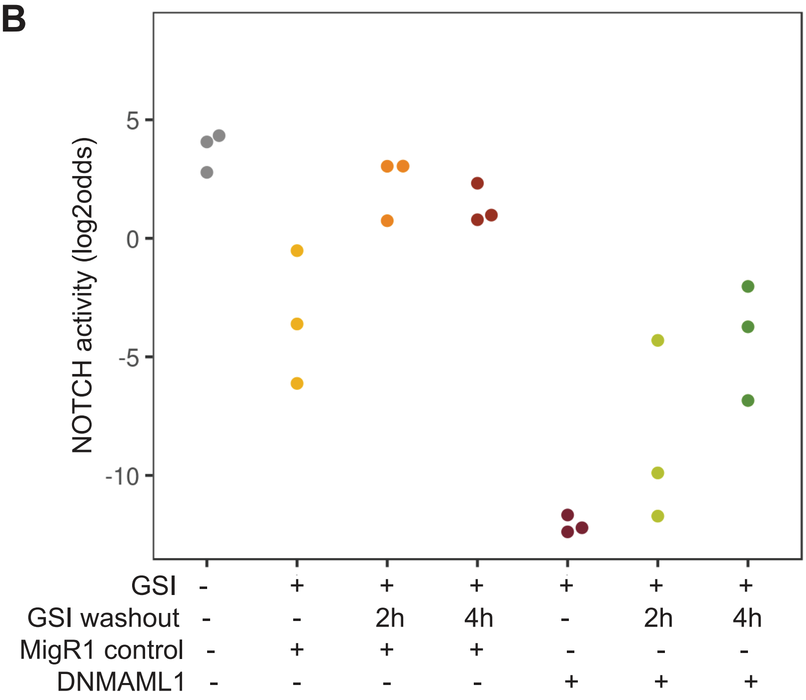

**Supplemental Figure S2. Correlations of ICN1 level and Notch pathway activity, divided per NOTCH1-activating mutation status.**

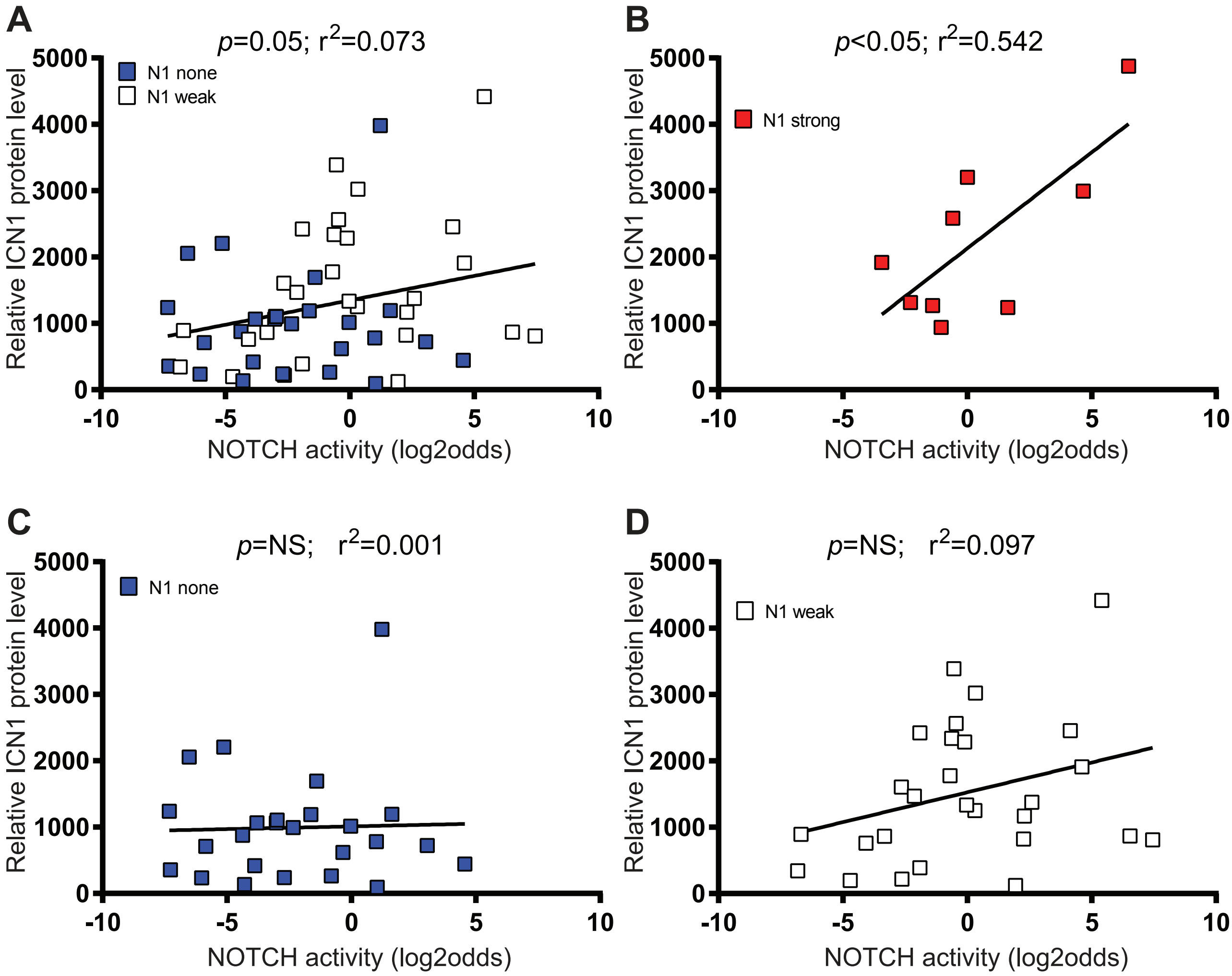
